# Plant-mycorrhizal associations may explain the latitudinal gradient of plant community assembly

**DOI:** 10.1101/2023.06.10.544476

**Authors:** Naoto Shinohara, Yuta Kobayashi, Keita Nishizawa, Kohmei Kadowaki, Akira Yamawo

## Abstract

Biogeographical variation in community assembly patterns forms the basis of large-scale biodiversity gradients. Previous studies on community assembly have suggested the dominant role of environmental filtering at higher latitudes. However, recent evidence indicates that plant community distributions at higher latitudes exhibit more spatial clustering, suggesting the importance of spatial assembly processes. In this study, we propose a hypothesis that resolves this paradox by incorporating biogeographic variations in dominant mycorrhizal types: the increasing prevalence of ectomycorrhizal (EcM) trees with latitude contributes to the spatially clustered distribution of plants, as EcM trees tend to exhibit positive plant-soil feedback. We analyzed a large-scale standardized dataset of Japanese forests covering a latitudinal gradient of >10° and found that (i) the proportion of EcM trees was higher at higher latitudes, and (ii) EcM tree-rich communities exhibited more spatially clustered distributions. Consequently, (iii) the tree species composition at higher latitudes was better explained by spatial variables. Moreover, consistent with predictions derived from the plant-soil feedback theory, these findings were particularly evident in the understory rather than the canopy communities. This study demonstrates that plant-soil feedback influences plant community distributions at metacommunity scales, thereby giving rise to a paradox within the latitudinal gradient of plant community assembly.

## Introduction

Understanding the ecological mechanisms that drive latitudinal biodiversity gradients has long been a challenge in the field of ecology (Pianka 1966; Hillebrand 2004). Classical studies have primarily focused on biogeographical variations in the mechanisms that maintain α-diversity (Lambers et al. 2002; Willig et al. 2003; Comita et al. 2014). However, recent attention has been directed towards elucidating the processes that generate β-diversity, which represents another crucial aspect of γ-diversity (Myers et al. 2013; Xing and He 2019; Nishizawa et al. 2022).β-diversity, which denotes a compositional variation in space, is a result of several community assembly processes. For instance, correlations between species composition and environmental variables are indicative of environmental filtering. In contrast, spatial correlations in β-diversity can reflect other spatial or stochastic processes, such as dispersal limitations or priority effects (Borcard et al. 1992; Tuomisto et al. 2003; Cottenie 2005; Shinohara et al. 2023).

Recent evidence suggests that the relative importance of processes driving β-diversity varies with latitude. In a pioneering study, Myers et al. (2013) suggested that environmental filtering is more prominent in temperate forests, whereas other spatial or stochastic processes dominate in tropical forests. The notion that the strength of environmental filtering increases with latitude aligns well with other community assembly studies; stronger environmental filtering is expected in habitats with higher abiotic stress (Chase 2007; Guo et al. 2014), lower productivity (Chase 2010; Kardol et al. 2013), and smaller species pool sizes (Fukami 2004), all of which characterize forests at higher latitudes. However, contradicting evidence challenges this paradigm. Qian and Ricklefs (2007) showed that climatic variables have minimal explanatory power in plant species composition in the northernmost part of North America. Moreover, a recent meta-analysis revealed that the explanatory power of environmental variables shows a hump-shaped response against latitude, indicating that the strength of environmental filtering peaks at mid-latitudes (approximately 30°) and decreases at higher latitudes (Nishizawa et al. 2022). However, the current paradigm of community assembly studies fails to account for the weaker effects of environmental filtering at higher latitudes. In this study, we propose a new hypothesis suggesting that biogeographical variations in plant-mycorrhizal associations modulate the latitudinal gradient in plant community assembly.

Feedback between plants and soil microbiota (plant-soil feedback, PSF), such as mycorrhizal fungi and soil pathogens, plays a critical role in plant community dynamics (Bever et al. 2010; Kadowaki et al. 2018; Ke and Wan 2020). Reciprocal interactions between plants and mycorrhizal fungi create positive PSF, whereas the accumulation of host-specific soil pathogens leads to negative PSF (Bever et al. 2010; van der Putten et al. 2013). Previous studies have suggested that the balance of positive and negative PSF, as well as the direction of net PSF, can be predicted based on the mycorrhizal types of plant species: trees associated with ectomycorrhizal (EcM) and arbuscular (AM) fungi tend to exhibit positive and negative PSF, respectively (Bennett et al. 2017; Kadowaki et al. 2018). One possible reason for the positive feedback observed in EcM trees is related to EcM fungi, which physically cover host roots with a fungal mantle to protect them from pathogen attack (Brundrett et al. 1990; Bennett et al. 2017) or produce extracellular enzymes that enhance the mineralization of complex organic substrates (Read et al. 2004; Liu et al. 2018).

While plant-soil feedback arises from local interactions between individual trees and soil microbiota, the direction of local-scale plant-soil feedback can have broader implications for the distribution of plant species within communities (Johnson et al. 2018; Sasaki et al. 2019). For instance, Johnson et al. (2018) demonstrated that saplings of EcM trees were more spatially clustered around adult EcM trees compared to AM saplings around AM adults, likely because of positive PSF. Consequently, the processes by which local-scale interactions contribute to the large-scale distribution of tree communities vary depending on the mycorrhizal tree species, potentially influencing β-diversity and its correlation with spatial patterns (Zhong et al. 2021). Notably, the large-scale consequences of PSF may drive the latitudinal gradient of plant community assembly because the type of mycorrhizal association varies with latitude; EcM trees are predominant at higher latitudes, whereas AM plants are more common at lower latitudes (Read et al. 2004; Barceló et al. 2019; Steidinger et al. 2019).

These findings collectively indicate that at higher latitudes, where EcM trees are more dominant, positive PSF is stronger, resulting in spatially clustered distributions of trees. This could explain why compositional variations in forests at higher latitudes are better explained by spatial variables rather than environmental variables (Qian and Ricklefs 2007; Nishizawa et al. 2022). We tested this hypothesis by analyzing a large dataset of Japanese forests. The elongated shape of the Japanese Archipelago (spanning from 25° N to 45° N) makes it an ideal system for investigating the latitudinal gradient of community assembly. Within this latitudinal range, a previous meta-analysis (Nishizawa et al. 2022) predicted an increase in the spatial clustering of plant community distributions with latitude. Using the dataset, we tested the following specific predictions: (i) a higher proportion of EcM trees at higher latitudes, (ii) greater spatial clustering in EcM tree-rich communities, and, consequently, (iii) a stronger influence of spatial variables on tree species compositions at higher latitudes. These predictions were examined separately for the forest canopy and understory. We further expected that (iv) the above predictions would be supported more clearly for understory communities than for canopy communities, as plant-soil feedback is more influential at earlier life stages (Chen et al. 2019; Sasaki et al. 2019).

## Material and Methods

### Data collection

We analyzed the national vegetation dataset obtained from the Ministry of the Environment of Japan (www.biodic.go.jp, accessed on April 15, 2021). The surveys were conducted from 2000 to 2019 using the phytosociological method, which involved recording coverage classes (labeled as 0.1, 1, 2, 3, 4, or 5, in order of increasing dominance) of plant species at each layer (labeled as 1 to 7, from the forest canopy to the moss layer) (Kobayashi et al. 2023). Our analyses focused solely on tree species because EcM mycorrhizal associations are typically reported for tree species (Soudzilovskaia et al. 2020).

A total of 37,276 plots were surveyed, but their sizes varied across surveys (median:105 m^2^; range:0.02–22,500 m^2^). To mitigate potential biases, we selected the data from plot with sizes of 100, 225, or 400 m^2^, as these were the most common sizes (4,992, 6,033, and 6,000 plots, respectively). Analyses were performed separately for the data obtained from different plot sizes. We excluded plots situated in non-forested areas. This data screening process involved creating a 100 m buffer around each plot and determining the proportion of land-use types (categorized as *forest*, *grassland*, *agricultural area*, or *others*) based on the high-resolution land use and land cover map (2014–2016, v.18.03) produced by the Japan Aerospace Exploration Agency, with a spatial resolution of 30 m (Hashimoto et al. 2014; https://www.eorc.jaxa.jp/ALOS/jp/dataset/lulc_j.htm). Plots were classified as forest if the most frequent land-use cell within the buffer was *forest*; otherwise, they were classified as non-forest. Some plots in alpine zones (situated at altitudes higher than 1000m) were also excluded from the analyses, to focus specifically on the latitudinal gradient and minimize the confounding effect of elevation. We further excluded plots presumably located in artificial forests by determining whether any of the typically planted trees in Japan (*Cryptomeria japonica*, *Chamaecyparis obtuse*, *Larix kaempferi*, and *Abies sachalinensis*) were dominant (i.e., the coverage class was 5).

For each plot, we obtained climatic and topographical variables using 1 km mesh climate data from the Land, Infrastructure, and Transportation Ministry (www.mlit.go.jp/en/index.html, accessed on June 7, 2015) and a digital elevation model with a spatial resolution of 30m from Yamazaki et al. (2020). We retrieved the following important environmental variables: (i) annual precipitation (mm), (ii) annual mean temperature (°C), (iii) elevation (m), and (iv) mean slope (°). The last two topographical variables in the list were calculated as the averages within the 100 m buffer around each plot.

For each tree species, the mycorrhizal association type was assigned based on an extensive database of plant mycorrhizal associations, FungalRoot (Soudzilovskaia et al. 2020). This database synthesizes information from published literature on mycorrhizal colonization worldwide and provides genus-level assignments of plant-mycorrhizal associations, which serve as reliable predictors. Following the authors’ recommendations, we assigned each species’ plant– mycorrhizal association based on its genus. The FungalRoot database indicated some cases of mixed colonization for a genus by two types of mycorrhizal fungi (“EcM-AM”) or facultative associations (“AM-NM”); we considered each species as either AM- or EcM-associated only if its mycorrhizal association could be reliably predicted from the genus, as indicated by “AM” or “EcM” in the FungalRoot database.

### Data analyses

First, the dataset was separated into two subsets based on the vertical forest layer. The first subset included species occurrences in the two highest-layer classes (classes 1 and 2), corresponding to the tallest and sub-tallest layers (referred to as the forest canopy data). The second subset included lower-layer classes 3 and 4, which mainly consisted of shrubs (referred to as the forest understory data).

The most common approach for quantifying the spatial clustering of community structures is variation partitioning (Borcard et al. 1992; Cottenie 2005; Shinohara et al. 2023). It partitions the compositional variation among sites into components explained by (i) spatial variables (R^2^_spa|env_), (ii) environmental variables (R^2^_env|spa_), and spatially correlated environmental variables (R^2^_both_). R^2^_spa|env_ represents the extent to which community compositions are spatially structured, independent of the distribution of background environments. When conducting variation partitioning, it is crucial to carefully define the metacommunity as a unit of analysis, as the choice of unit size may bias the inference (Viana and Chase 2019; Nishizawa et al. 2022).

Thus, we aimed to be as non-arbitrary as possible in selecting the unit. In this study, we defined the standard second mesh as the unit of metacommunities, as prescribed and used in various Japanese government statistics. The second mesh divides Japanese areas based on coordinates by 5’ in the latitudinal and 7’ 30’ in the longitudinal directions, resulting in each mesh covering an area of approximately 100 km^2^. This metacommunity size (a unit of the variation partitioning analysis) aligns with in the common practice in many plant community assembly studies (51.6% percentile among 171 studies collected by Nishizawa et al. 2022). To ensure statistically reliable results for the variation partitioning analysis, we restricted our analyses to metacommunities that included 10 plots or more per plot size (note that the data obtained for different plot sizes [100, 225, and 400 m^2^] were analyzed separately). As a result, we had 48, 61, and 38 metacommunities (i.e., second meshes) for 100, 225, and 400 m^2^ plot sizes, respectively, for the forest canopy data, and 56, 63, and 44 metacommunities for 100, 225, and 400 m^2^ plot sizes, respectively, for the forest understory data.

For each metacommunity, we conducted a variation partitioning analysis using a distance-based redundancy analysis (db-RDA, Legendre and Anderson 1999). db-RDAs were performed based on the Jaccard dissimilarity matrix, calculated as *S_ij_* / (*S_i_* + *S_j_* − *S_ij_*), where *S_ij_* is the number of species present in both plots *i* and *j* and *S_i_* and *S_j_* are the total number of species present in plots *i* and *j*, respectively. We implemented the db-RDA using a *capscale* function. The explanatory variables were divided into two classes: environmental and spatial variables. The environmental variables included climatic (annual precipitation and mean temperature) and topographic (elevation and average slope) conditions of each plot. We did not include quadratic terms of these variables in our modeling as has often been done to model unimodal responses of species to the environment, since their inclusion led to collinearity issues among explanatory variables. For this decision, we assumed that the ranges of environments in each metacommunity (ca. 100 km^2^ in area) are not so vast that species responses can be adequately modeled by linear relationships. Spatial variables were generated using the principal components of neighbor matrices (PCNM, Borcard and Legendre 2002) based on the coordinates (longitude and latitude) of the plots within each metacommunity, using the *pcnm* function. Eigenvectors with positive eigenvalues were used as explanatory variables. Before conducting the variation partitioning, variable selection was performed separately for environmental and spatial variables using the forward-backward stepwise method based on *p-*values. The median number of selected environmental variables was 1 (range: 0–4) and that of the spatial variables was 2 (range: 0–8). Variable selection was performed using the *ordistep* function. Subsequently, the selected sets of environmental and spatial variables were subjected to variation partitioning analysis using the *varpart* function. The returned R^2^ values were adjusted for the number of predictors and sample size using the *RsquareAdj* function (Peres-Neto et al. 2006). To meet the distributional condition for the linear regression models (specified later), the adjusted R^2^ values were set to zero if they were negative or if no spatial variables were selected in the variable selection.

We constructed generalized linear models (GLMs) to examine the relationship between (i) latitude and the proportion of EcM trees at the metacommunity (second-mesh) level, (ii) the proportion of EcM trees and the explanatory power of spatial variables for compositional variations (R^2^_spa|env_), and (iii) latitude and R^2^_spa|env_. The first GLM was modeled with a binomial distribution (logit link) included data from all second meshes data (including those with fewer than 10 plots). The second and third GLMs were modeled with a quasi-binomial distribution (logit link) because the R^2^_spa|env_ values were concentrated around zero (underdispersed). The GLMs were constructed separately for different plot sizes (100, 225, and 400 m^2^) and for the forest canopy and understory layers, resulting in six combinations per model.

All data screening and statistical procedures were performed using ARCGIS PRO version 2.6.0 (ESRI) and R version 4.1.3 (R Core Team 2022) using the *vegan* package (Oksanen et al. 2022). Figures were generated using the *ggplot2* package (Wickham 2016).

## Results

The metacommunities that met our criterion for analysis (i.e., including ≥10 plots) were distributed across a range of latitudes (approximately 8°, 7°, and >12° for data with plot sizes of 100, 225, and 400 m^2^, respectively; Appendix 1: Fig. S1). In favor of covering the widest latitudinal range, we have presented the results for the 400 m^2^ plot size data in the main text. The results for the 100 and 225 m^2^ plots are presented in the *Supporting Information* (Appendix 1: Figs. S2, S3, and S4; Tables S1, S2, and S3).

A total of 482 and 683 tree species were recorded in the forest canopy and understory layers, respectively. AM trees were dominant in both datasets, with 349 species in the canopy dataset and 524 species in the understory dataset. The number of EcM trees was lower, with 77 species in the canopy dataset and 58 species in the understory dataset. The proportion of EcM trees varied considerably across metacommunities, ranging from 4.6–100% in forest canopy and from 2.0–100% in the understory (Fig. 1). Notably, the proportion of EcM trees was higher at higher latitudes in both the forest canopy (slope: 0.039; *p* < 0.001; Fig. 1, Appendix 1: Table S1) and understory (slope: 0.056, *p* < 0.001).

**Figure 1.**
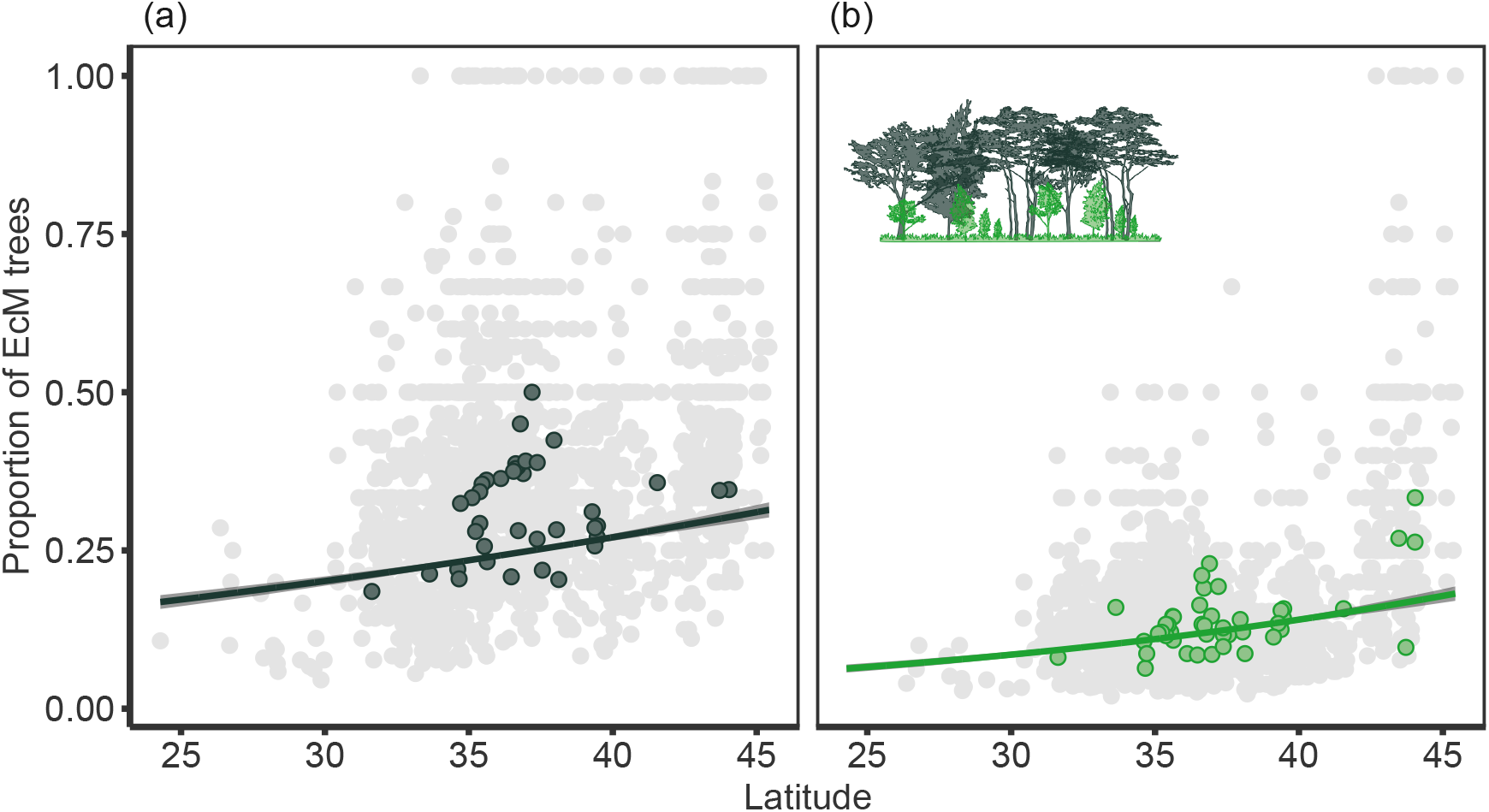
Relationships between the proportion of EcM trees within a second mesh (a unit of metacommunity) in (a) the canopy and (b) understory layers, and latitude for data obtained with a 400 m^2^ plot size. The colored points represent the second meshes that met our criteria and were used in the other analyses. The lines and the 95% confidence intervals are drawn based on linear regression.

For the forest understory data, the explanatory power of the spatial variables for compositional variation (R^2^_spa|env_) was positively associated with the proportion of EcM trees (slope: 6.50; *p* = 0.007, Fig. 2, Appendix 1: Table S2). A significant relationship between these variables was not observed for the forest canopy (slope: 0.48; *p* = 0.827).

**Figure 2.**
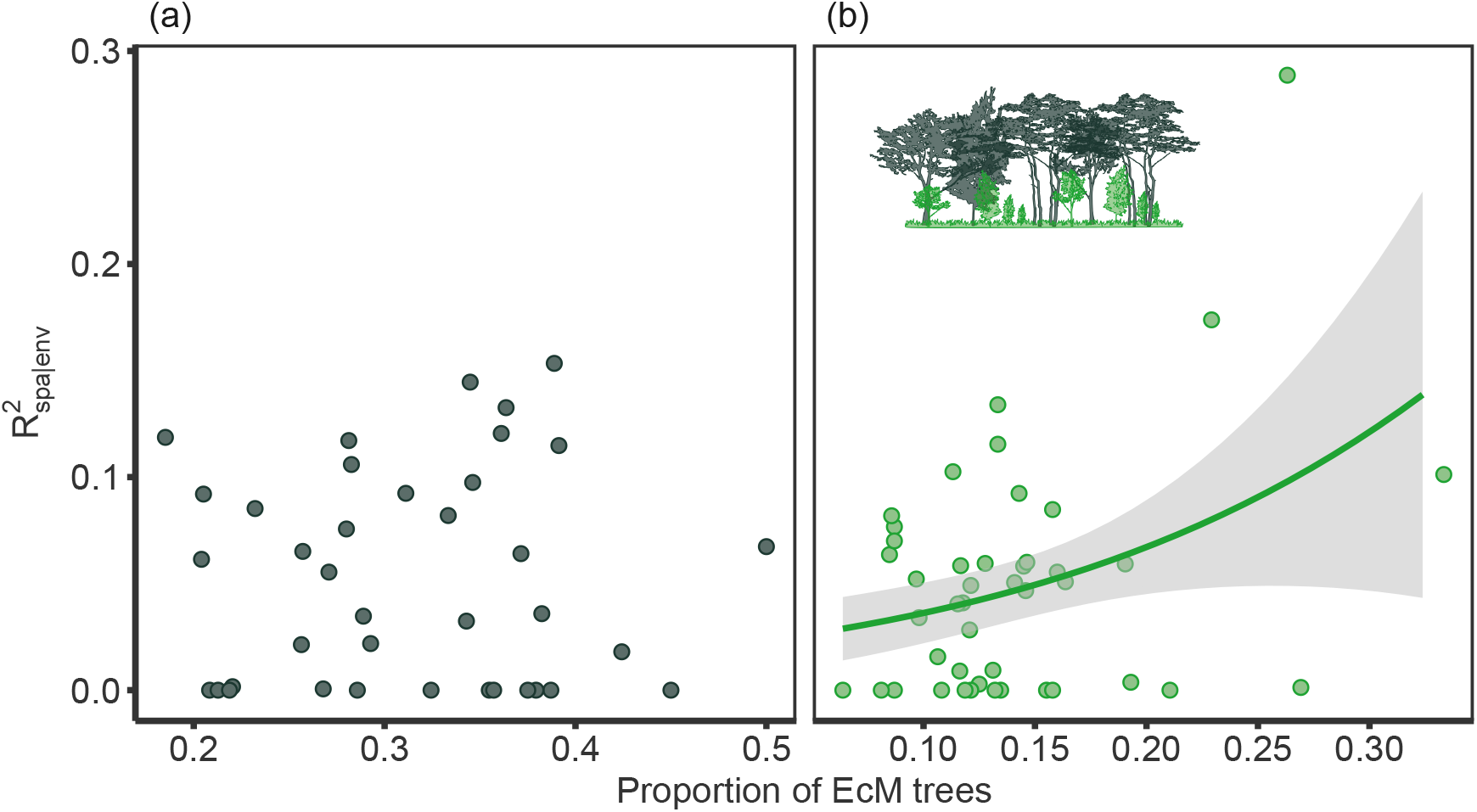
The metacommunity-level relationships of the proportion of EcM trees with the explanatory power of spatial variables for among-plots compositional variation of tree communities (R^2^_spa|env_) for the (a) canopy and (b) understory layers. The line and the 95% confidence interval are drawn based on linear regression if the slope significantly deviated from zero (*p* &lt; 0.05).

The results also showed that the R^2^_spa|env_ value increased with an increase in latitude in the forest understory data (slope: 0.127; *p* = 0.020; Fig. 3, Appendix 1: Table S3). Conversely, this relationship was not statistically significant for the forest canopy (slope: 0.053; *p* = 0.41; Fig. 2, Appendix 1: Table S2).

**Figure 3.**
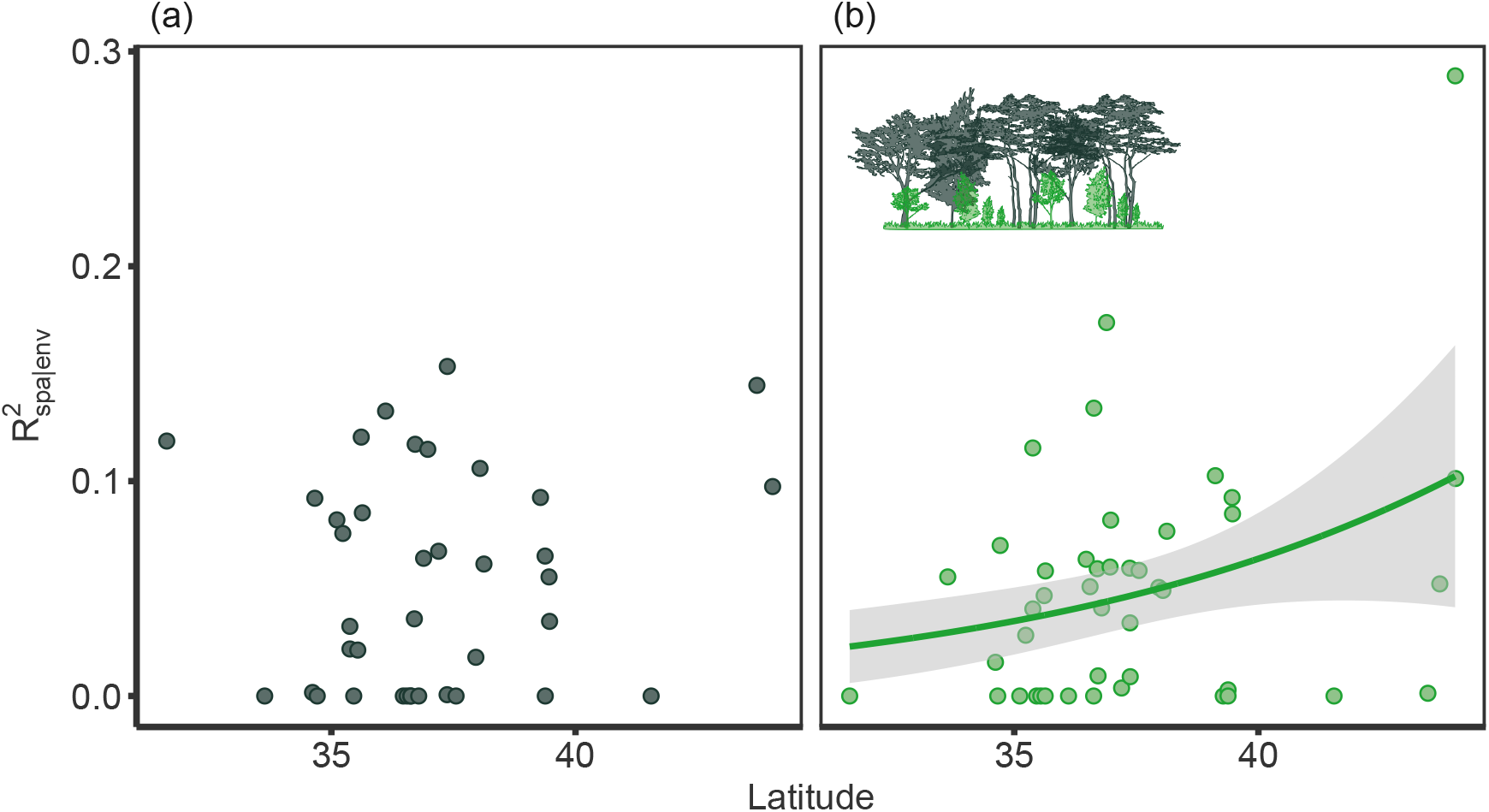
The metacommunity-level relationships between latitude and the explanatory power of spatial variables for among-plots compositional variation of tree communities (R^2^_spa|env_) in the (a) canopy and (b) understory layers. The line and the 95% confidence interval are drawn based on linear regression if the slope significantly deviated from zero (*p* &lt; 0.05).

## Discussion

In this study, we have demonstrated a significant association between the proportion of EcM trees and the spatial structure of community composition at the metacommunity level in the forest understory. Our findings reveal that communities dominated by EcM trees exhibit greater spatial clustering (Fig. 2b), likely reflecting the stronger PSF exhibited by EcM tree species compared to AM tree species (Bennett et al. 2017; Kadowaki et al. 2018). The findings are largely unaffected by the potential confounding effects of environmental variables associated with latitude, since the variation partitioning analysis allowed us to disentangle purely spatial processes from processes driven by environmental heterogeneity, including those correlated with space.

A common observation in studies on β-diversity is the distance-decaying pattern, wherein species compositions are more similar between pairs of sites that are closer to each other (Condit et al. 2002; Soininen et al. 2007). This pattern of distance decay in compositional similarity has traditionally been attributed to various community assembly processes, including species sorting based on background abiotic environmental factors (Tuomisto et al. 2003) and limited dispersal capabilities of individuals (Clark et al. 1999; Condit et al. 2002). Our findings suggest that the distance decay of β-diversity can also be influenced by tree–mycorrhizal associations. Thus, this study provides a novel perspective in plant community assembly research, highlighting the potential significance of individual-level plant–mycorrhizal associations in shaping large-scale community patterns.

As anticipated, the correlation between the proportion of EcM trees and the degree of spatial clustering in local communities was observed solely in the forest understory layer (Fig. 2a) and not in the forest canopy. This outcome aligns with the prevailing perspective in PSF studies, which suggests that feedback effects exert a stronger influence during the early life stages of plants (Chen et al. 2019; Sasaki et al. 2019). However, as individuals mature, EcM seedlings in close proximity to adult trees experience stronger intraspecific competition (Hoeksema 2005). This competitive ‘self-thinning’ process could weaken the spatial aggregation of EcM saplings, thereby explaining why a high proportion of EcM trees was not associated with spatial clustering of species compositions in the canopy layer. The distribution patterns of established adult trees can also be affected by other factors, such as major disturbances (e.g., hurricanes or pest outbreaks) that may override the influence of plant-soil feedback experienced during earlier life stages.

Importantly, our findings provide an explanation for the spatially structured patterns of tree metacommunities at higher latitudes (Qian and Ricklefs 2007; Nishizawa et al. 2022), contrary to the prediction of the current paradigm in community assembly studies, which posits that environmental filtering would be stronger at higher latitudes (Fukami 2004; Chase 2007; Chase 2010; Kardol et al. 2013; Guo et al. 2014). The transition from AM-dominated to EcM-dominated forests with increasing latitude (Fig. 1) strongly suggests a shift in the direction of PSF shifting from negative to positive, resulting in more spatially clustered distributions of understory communities at higher latitudes (Fig. 3). Although previous studies have explored the possibility of PSF explaining the latitudinal gradient of tree α-diversity (Lambers et al. 2002; Comita et al. 2014), our findings complement them by proposing that plant-soil feedback and its dependence on mycorrhizal type may affect tree β-diversity and potentially mediate regional tree coexistence. Our results highlight the underappreciated role of tree-mycorrhizal associations in characterizing the determinants of β-diversity along a latitudinal gradient.

We found no significant correlations for the subsets of data obtained with small survey plot sizes (100 m^2^ or 225 m^2^; Appendix 1: Fig. S3, S4). There are two main reasons for the lack of clear patterns. First, during data processing, we removed data points that did not meet our criteria. Consequently, the latitudinal range covered in this study differed considerably among data with different plot sizes, ranging from > 12° for the 400 m^2^ plot size data to approximately 8° and 7° for the 100 and 225 m^2^ plot sizes, respectively (Appendix 1: Fig. S1). The narrower geographical ranges resulted in smaller variations in the proportion of EcM tree for the 100 and 225 m^2^ plot size data compared with the 400 m^2^ plot size data (compare the ranges of the x-axis for the forest understory subsets in Figs. 2 and Appendix 1: Fig. S3). These smaller variations in the explanatory variables may have reduced the statistical power to detect possible links. Alternatively, when surveys are conducted with smaller plot sizes, the effects of demographic stochasticity or small-scale random seed dispersal on species composition can be more pronounced (Gilbert and Levine 2017). Thus, at smaller plot sizes (100 and 225 m^2^), the influence of PSF on species composition could be obscured by the stochastic nature of community assembly.

Although our results are consistent with the hypothesis that PSF explain the latitudinal gradient of tree community assembly, other non-mutually exclusive explanations exist. A recent study suggested that EcM trees have lower seed dispersal ability compared to AM trees, likely because of the joint evolution of mycorrhizal type and dispersal ability (Yamawo and Ohno 2021). The shorter dispersal range of EcM trees could result in more spatially concentrated distributions, which could explain our findings (Fig. 2). Conversely, the long-distance dispersal ability of AM trees could limit their spatial clustering. Indeed, we found a weak negative correlation between the R^2^_spa|env_ of forest understory communities and the proportion of AM trees (GLM with quasi-binomial distribution, slope: −3.08; *p* = 0.30; Appendix 1: Fig. S5), suggesting that the forests dominated by AM trees were less spatially clustered. Given the limitations of our correlative approach, it remains a subject for future follow-up studies to investigate whether the spatially clustered distribution in forests dominated by EcM trees reflects the effect of positive PSF or the shorter dispersal ability of EcM trees. Further tests conducted in other regions are also necessary to validate the generalizability of our findings. Nevertheless, we emphasize that our results are consistent with our hypothesis, leading us to conclude that plant-mycorrhizal associations may explain the latitudinal gradient of plant community assemblies.

## Acknowledgments

This study was supported by Grant-in-Aid for JSPS Fellows 23KJ0077 (to NS) and a Grant-in-Aid for Scientific Research 21H02233 (to KK).

## Appendix 1

**Figure. S1.**
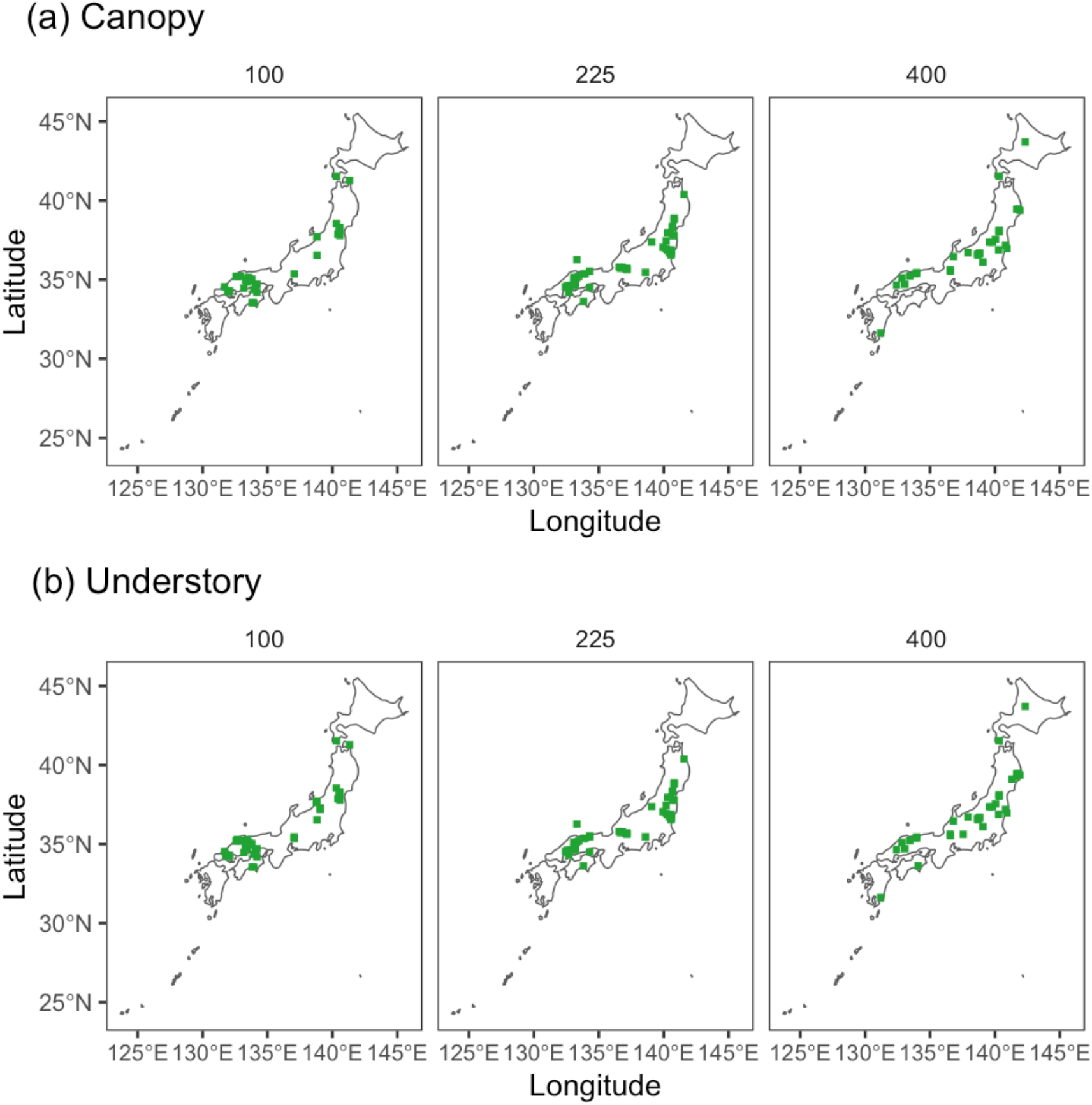
The distributions of second meshes (or metacommunities) that met our criterion and used in the analyses for (a) forest canopy and (b) understory layers. Note that the analyses were performed separately for the data obtained in different plot sizes (100, 225, and 400 m^2^) and thus the maps are presented separately.

**Figure. S2.**
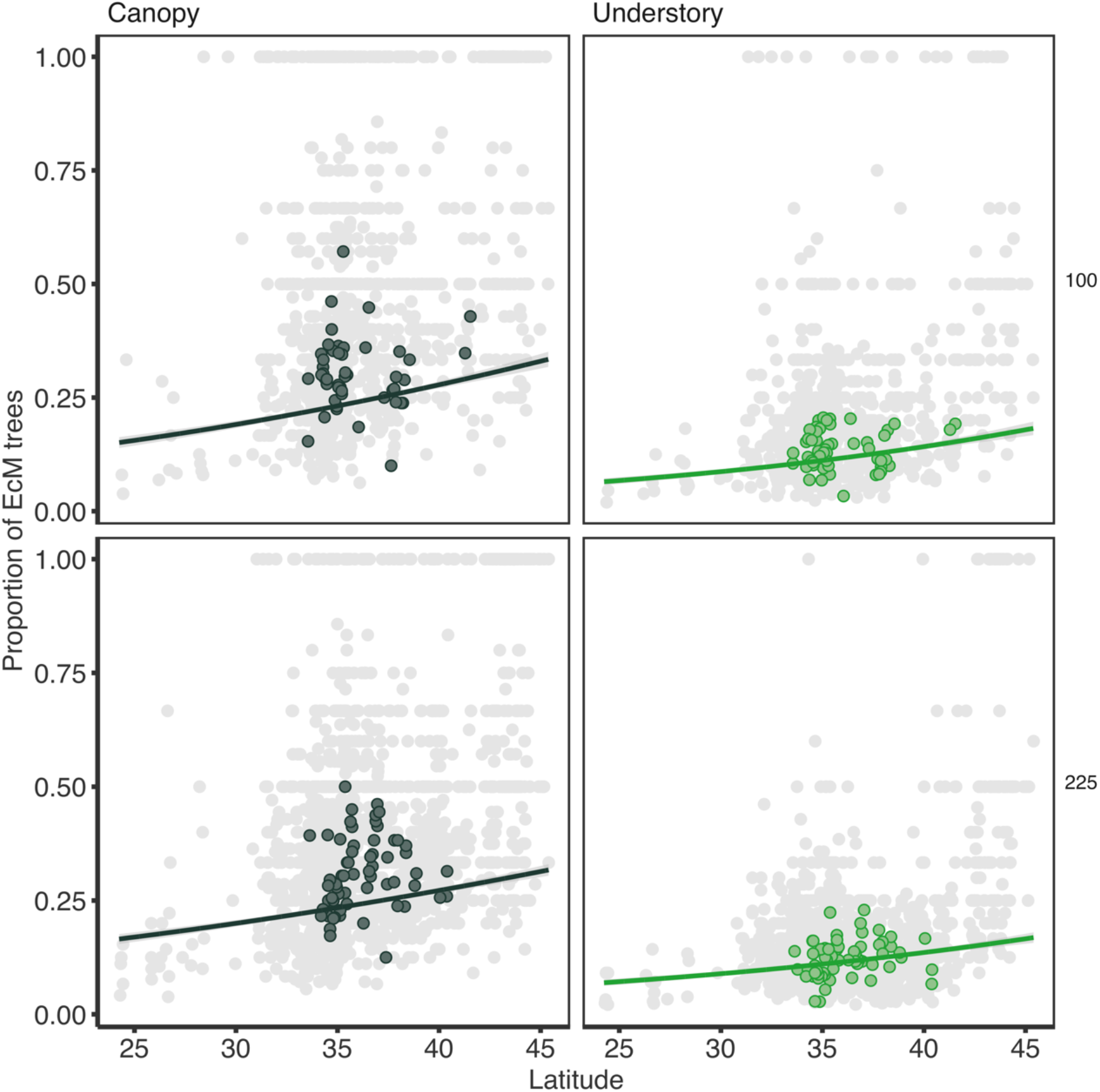
Relationships between the proportion of EcM trees within a second-mesh (a unit of metacommunity) in the canopy (left column) and understory (right column) layers, and latitude. The relationships are separately shown for data obtained with 100 (upper panels) and 225 m^2^ (lower panels) plot size. The colored points represent the second mesh that met our criterion and were used in the other analyses. The lines and the 95 % confidence intervals are drawn based on linear regressions.

**Figure. S3.**
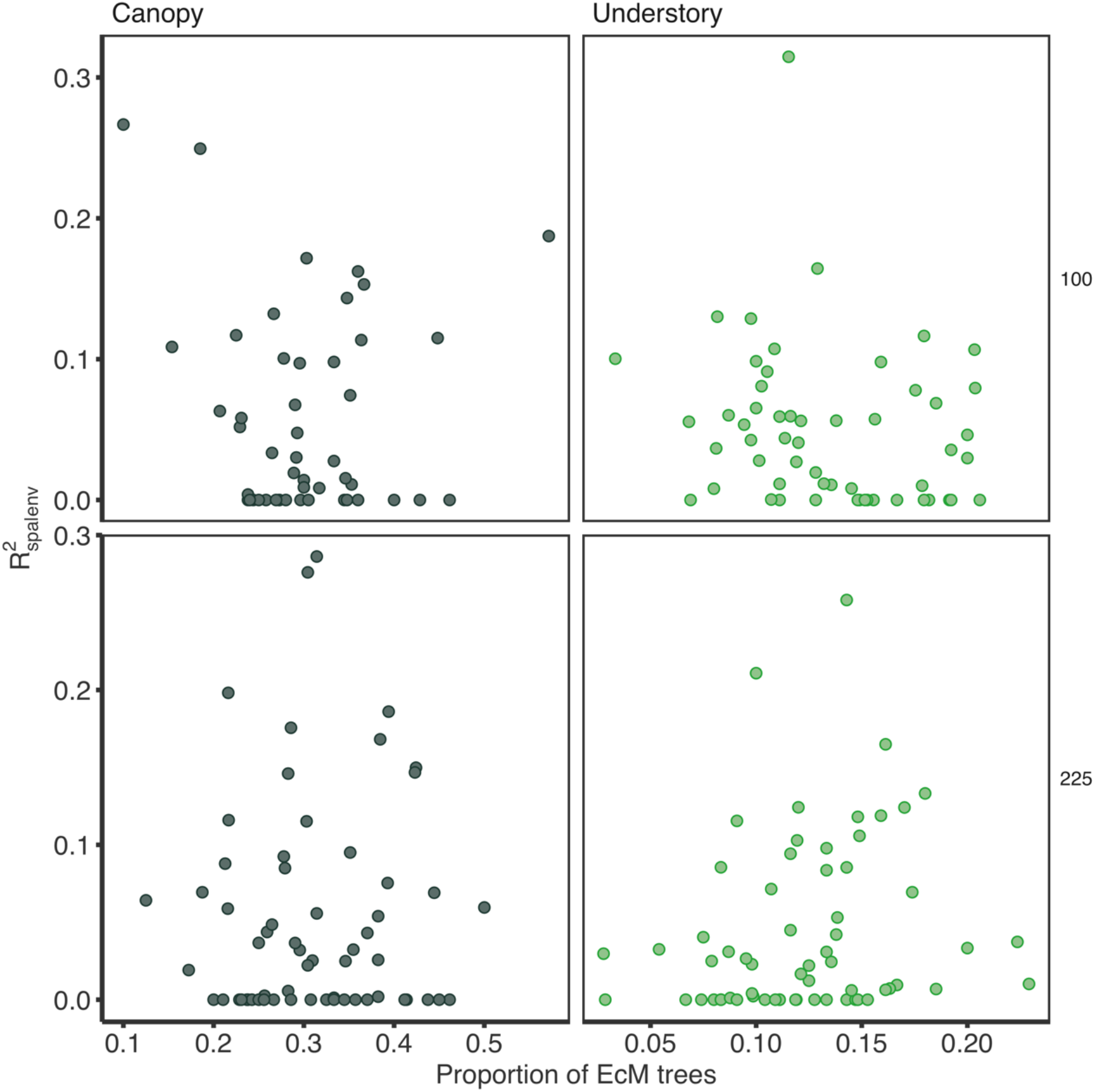
The metacommunity-level relationships of the proportion of EcM trees with the explanatory power of spatial variables for among-plots compositional variation of tree communities (R^2^_spa|env_) for the canopy (left columns) and understory (right columns) layers. The relationships are separately shown for data obtained with 100 (upper panels) and 225 m^2^ (lower panels) plot size. None of the relationships were statistically significant (*p* > 0.05, see table S2).

**Figure. S4.**
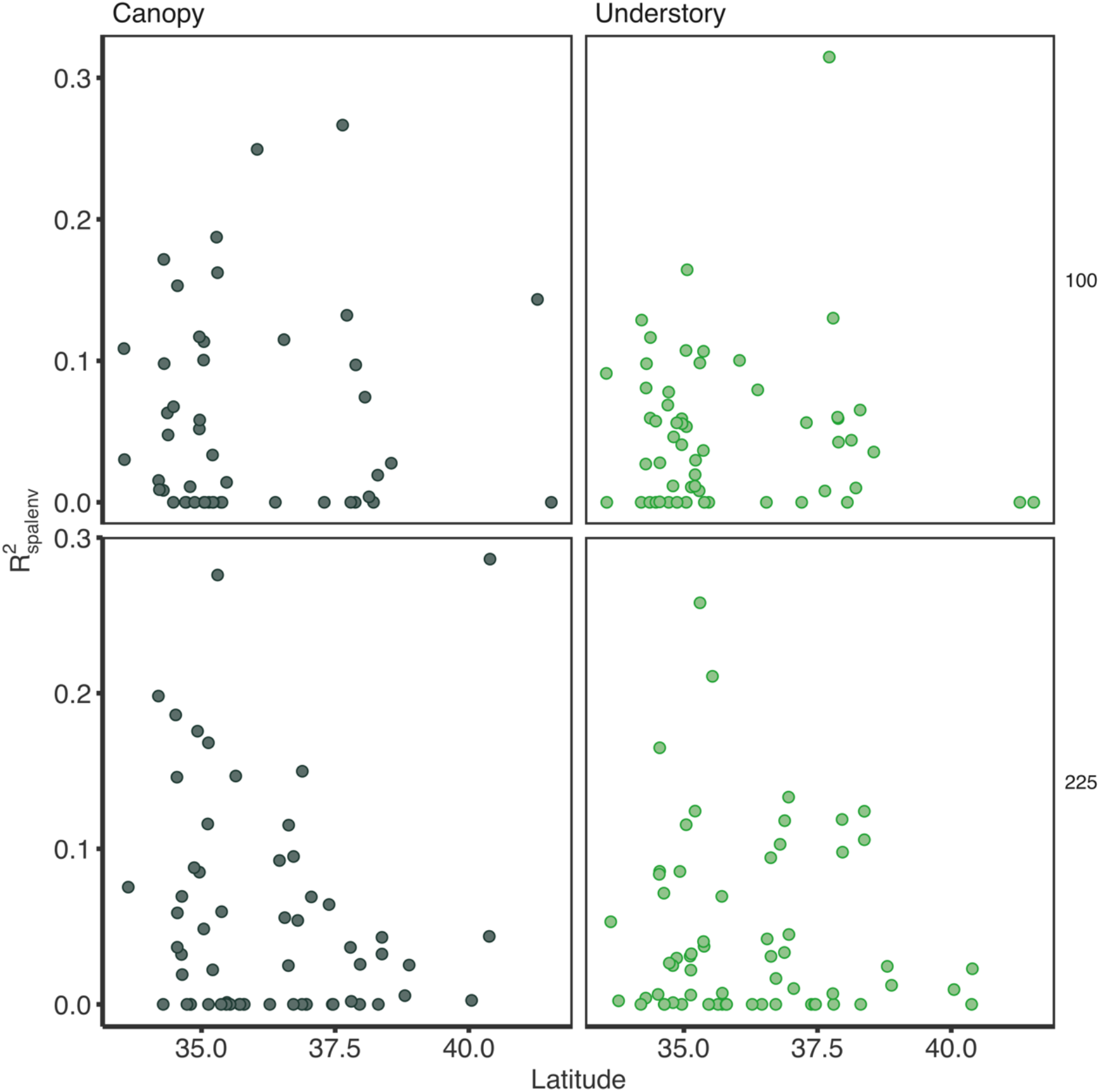
The metacommunity-level relationships between latitude and the explanatory power of spatial variables for among-plots compositional variation of tree communities (R^2^_spa|env_) in the canopy (left columns) and understory (right columns) layers. The relationships are separately shown for data obtained with 100 (upper panels) and 225 m^2^ (lower panels) plot size. None of the relationships were statistically significant (*p* > 0.05, see table S3).

**Figure. S5.**
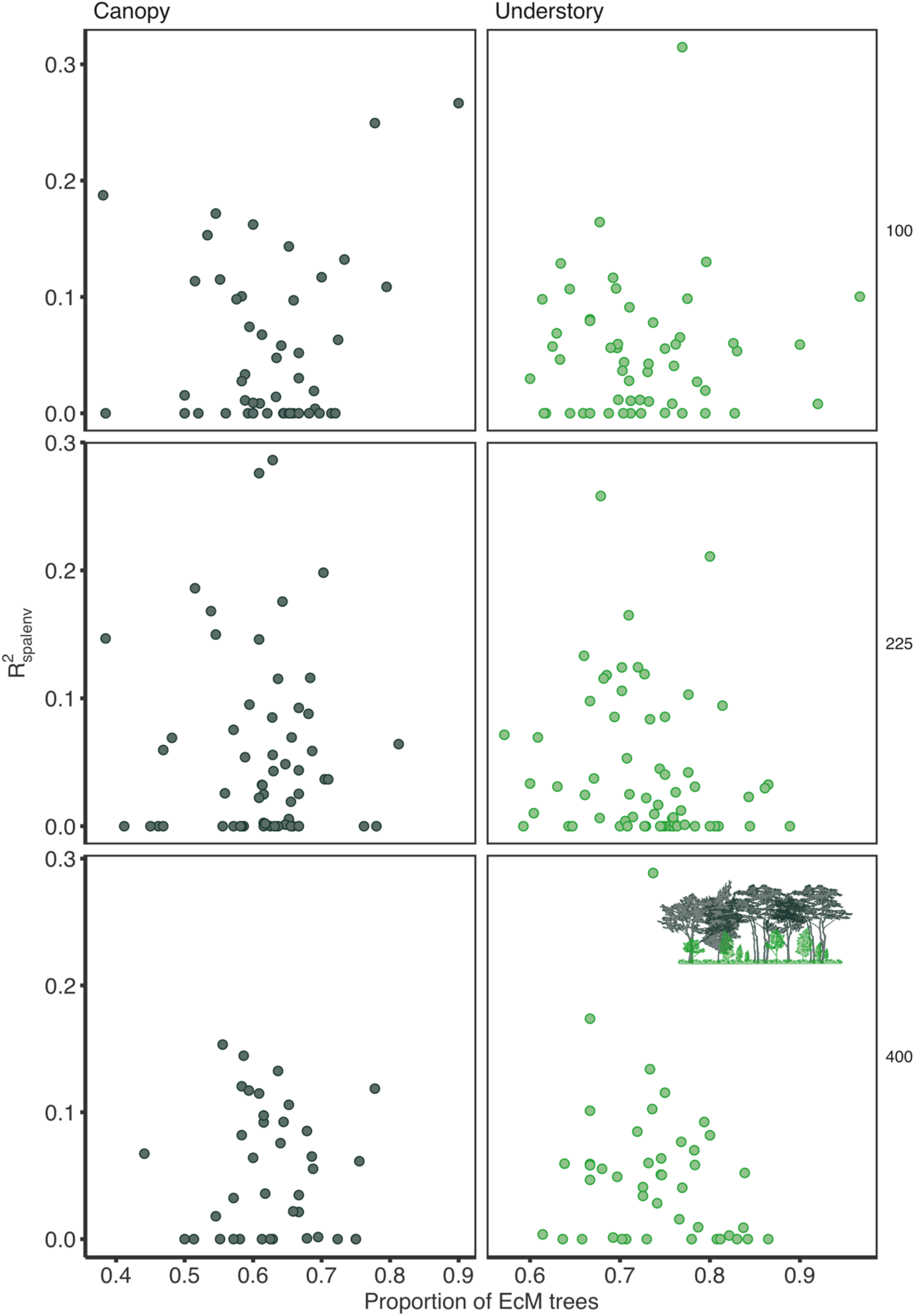
The metacommunity-level relationships of the proportion of AM trees with the explanatory power of spatial variables for among-plots compositional variation of tree communities (R^2^_spa|env_) for the canopy (left columns) and understory (right columns) layers. The relationships are separately shown for data obtained with 100 (upper row), 225 (middle row), and 400 m^2^ (lower row) plot size. None of the relationships were statistically significant (GLM with quasibinomial distribution, *p* > 0.05).

**Table S1.** Summary of the generalized linear models where the response variable was the proportion of EcM trees within a second-mesh (a unit of metacommunity) and the explanatory was latitude (see Fig. 1 and S2). The estimates of the slope and corresponding standard errors (SEs) and *p*-values are shown.

| Plot size | Canopy |  |  | Floor |  |  |
| --- | --- | --- | --- | --- | --- | --- |
|  | Estimate | SE | *p* | Estimate | SE | *p* |
| 100 | 0.0489 | 0.0039 | *p*<0.001 | 0.0549 | 0.0052 | *p*<0.001 |
| 225 | 0.0403 | 0.0032 | *p*<0.001 | 0.0474 | 0.0045 | *p*<0.001 |
| 400 | 0.0385 | 0.0030 | *p*<0.001 | 0.0562 | 0.0042 | *p*<0.001 |

**Table S2.**
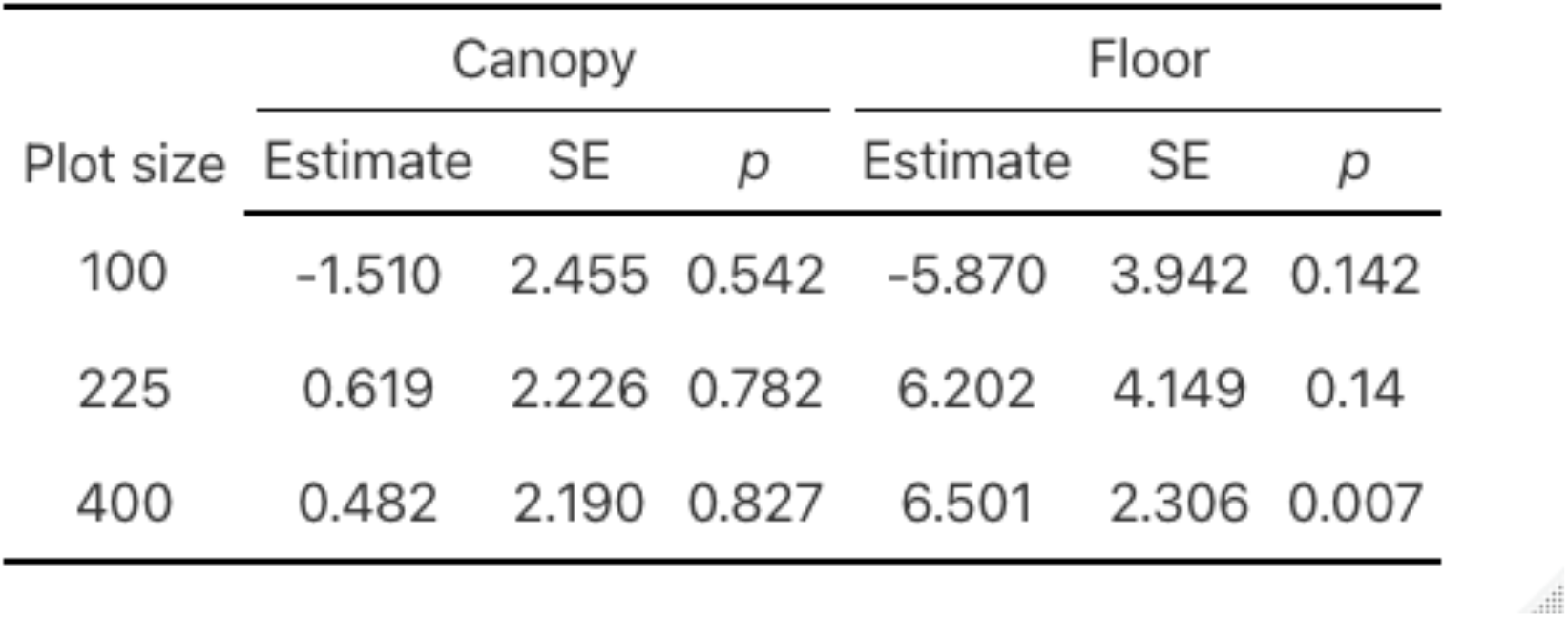
Summary of the generalized linear models where the response variable was the explanatory power of spatial variables for among-plots compositional variation of tree communities within a metacommunity (adjusted R^2^_spa|env_) and the explanatory was the proportion of EcM trees (see Fig. 2 and S3). The estimates of the slope and corresponding standard errors (SEs) and *p*-values are shown.

| Plot size | Canopy |  |  | Floor |  |  |
| --- | --- | --- | --- | --- | --- | --- |
| | Estimate | SE | $p$ | Estimate | SE | $p$ |
| 100 | -1.510 | 2.455 | 0.542 | -5.870 | 3.942 | 0.142 |
| 225 | 0.619 | 2.226 | 0.782 | 6.202 | 4.149 | 0.14 |
| 400 | 0.482 | 2.190 | 0.827 | 6.501 | 2.306 | 0.007 |

**Table S3.** Summary of the generalized linear models where the response variable was the explanatory power of spatial variables for among-plots compositional variation of tree communities within a metacommunity (adjusted R^2^_spa|env_) and the explanatory was latitude (see Fig. 3 and S4). The estimates of the slope and corresponding standard errors (SEs) and *p*-values are shown.

| Plot size | Canopy |  |  | Floor |  |  |
| --- | --- | --- | --- | --- | --- | --- |
| | Estimate | SE | $p$ | Estimate | SE | $p$ |
| 100 | 0.025 | 0.102 | 0.807 | -0.020 | 0.094 | 0.834 |
| 225 | -0.092 | 0.122 | 0.456 | -0.051 | 0.111 | 0.652 |
| 400 | 0.053 | 0.064 | 0.41 | 0.127 | 0.052 | 0.02 |

## References

Barceló M, Bodegom PM, Soudzilovskaia NA (2019) “Climate drives the spatial distribution of mycorrhizal host plants in terrestrial ecosystems.” J Ecol 107 (6):2564–2573

Bennett JA, Maherali H, Reinhart KO, Lekberg Y, Hart MM, Klironomos J (2017) “Plant-soil feedbacks and mycorrhizal type influence temperate forest population dynamics.” Science 355 (6321):181–184

Bever JD, Dickie IA, Facelli E, Facelli JM, Klironomos J, Moora M, Rillig MC, Stock WD, Tibbett M, Zobel M (2010) “Rooting theories of plant community ecology in microbial interactions.” Trends Ecol Evol 25 (8):468–478

Borcard D, Legendre P (2002) “All-scale spatial analysis of ecological data by means of principal coordinates of neighbour matrices.” Ecol Modell 153 (1–2):51–68

Borcard D, Legendre P, Drapeau P (1992) “Partialling out the spatial component of ecological variation.” Ecology 73 (3):1045–1055

Brundrett M, Murase G, Kendrick B (1990) “Comparative anatomy of roots and mycorrhizae of common Ontario trees.” Can J Bot 68 (3):551–578

Chase JM (2007) “Drought mediates the importance of stochastic community assembly.” Proc Natl Acad Sci U S A 104 (44):17430–17434

Chase JM (2010) “Stochastic community assembly causes higher biodiversity in more productive environments.” Science 328 (5984):1388–1391

Chen L, Swenson NG, Ji N, Mi X, Ren H, Guo L, Ma K (2019) “Differential soil fungus accumulation and density dependence of trees in a subtropical forest.” Science 366 (6461):124–128

Clark JS, Silman M, Kern R, Macklin E, HilleRisLambers J (1999) “Seed dispersal near and far: Patterns across temperate and tropical forests.” Ecology 80 (5):1475–1494

Comita LS, Queenborough SA, Murphy SJ, Eck JL, Xu K, Krishnadas M, Beckman N, Zhu Y, Gómez-Aparicio L (2014) “Testing predictions of the Janzen– Connell hypothesis: A meta-analysis of experimental evidence for distance- and density-dependent seed and seedling survival.” J Ecol 102 (4):845–856

Condit R, Pitman N, Leigh EG, Chave J, Terborgh J, Foster RB, Núñez P, Aguilar S, Valencia R, Villa G, Muller-Landau HC, Losos E, Hubbell SP (2002) “Beta-diversity in tropical forest trees.” Science 295 (5555):666–669

Cottenie K (2005) “Integrating environmental and spatial processes in ecological community dynamics.” Ecol Lett 8 (11):1175–1182

Fukami T (2004) “Community assembly along a species pool gradient: Implications for multiple-scale patterns of species diversity.” Popul Ecol 46 (2):137–147

Gilbert B, Levine JM (2017) “Ecological drift and the distribution of species diversity.” Proc Biol Sci 284 (1855):20170507

Guo H, Więski K, Lan Z, Pennings SC (2014) “Relative influence of deterministic processes on structuring marsh plant communities varies across an abiotic gradient.” Oikos 123 (2):173– 178

Hashimoto S, Tadono T, Onosato M, Hori M, Shiomi K (2014) “A new method to derive precise land-use and land-cover maps using multi-temporal optical data.” J Remote Sens Soc Jpn 34 (2):102–112

Hillebrand H (2004) “On the generality of the latitudinal diversity gradient.” Am Nat 163 (2):192–211

Hoeksema JD (2005) “Plant–plant interactions vary with different mycorrhizal fungus species.” Biol Lett 1 (4):439–442

Johnson DJ, Clay K, Phillips RP (2018) “Mycorrhizal associations and the spatial structure of an old-growth forest community.” Oecologia 186 (1):195–204

Kadowaki K, Yamamoto S, Sato H, Tanabe AS, Hidaka A, Toju H (2018) “Mycorrhizal fungi mediate the direction and strength of plant-soil feedbacks differently between arbuscular mycorrhizal and ectomycorrhizal communities.” Commun Biol 1 (1):196

Kardol P, Souza L, Classen AT (2013) “Resource availability mediates the importance of priority effects in plant community assembly and ecosystem function.” Oikos 122 (1):84–94

Ke P-J, Wan J (2020) “Effects of soil microbes on plant competition: A perspective from modern coexistence theory.” Ecol Monogr 90 (1):e01391

Kobayashi Y, Haga C, Shinohara N, Nishizawa K, Mori AS (2023) “Dominant temperate and subalpine Japanese trees have variable photosynthetic thermal optima according to site mean annual temperature.” Glob Ecol Biogeogr 32 (3):397–407

Lambers JH, Clark JS, Beckage B (2002) “Density-dependent mortality and the latitudinal gradient in species diversity.” Nature 417 (6890):732–735

Legendre P, Anderson MJ (1999) “DISTANCE-BASED REDUNDANCY ANALYSIS: TESTING MULTISPECIES RESPONSES IN MULTIFACTORIAL ECOLOGICAL EXPERIMENTS.” Ecol Monogr 69 (1): 1–24.

Liu X, Burslem DFRP, Taylor JD, Taylor AFS, Khoo E, Majalap-Lee N, Helgason T, Johnson D (2018) “Partitioning of soil phosphorus among arbuscular and ectomycorrhizal trees in tropical and subtropical forests.” Ecol Lett 21 (5):713–723

Myers JA, Chase JM, Jiménez I, Jørgensen PM, Araujo-Murakami A, Paniagua-Zambrana N, Seidel R (2013) “Beta-diversity in temperate and tropical forests reflects dissimilarmechanisms of community assembly.” Ecol Lett 16 (2):151–157

Nishizawa K, Shinohara N, Cadotte MW, Mori AS (2022) “The latitudinal gradient in plant community assembly processes: A meta-analysis.” Ecol Lett 25 (7):1711–1724

Oksanen J, Simpson GL, Blanchet FG, Kindt R, Legendre P, Minchin PR, O’Hara RB, et al. (2022) vegan: Community Ecology Package. R package version 2.6–2. https://CRAN.R-project.org/package=vegan

Peres-Neto PR, Legendre P, Dray S, Borcard D (2006) “Variation partitioning of species data matrices: Estimation and comparison of fractions.” Ecology 87 (10):2614–2625

Pianka ER (1966) “Latitudinal gradients in species diversity: A review of concepts.” Am Nat 100 (910):33–46

Qian H, Ricklefs RE (2007) “A latitudinal gradient in large-scale beta diversity for vascular plants in North America.” Ecol Lett 10 (8):737–744

R Core Team (2022). R: A Language and Environment for Statistical Computing. URL https://www.R-project.org/ R Foundation for Statistical Computing, Vienna, Austria

Read DJ, Leake JR, Perez-Moreno J (2004) “Mycorrhizal fungi as drivers of ecosystem processes in heathland and boreal forest biomes.” Can J Bot 82 (8):1243–1263

Sasaki T, Konno M, Hasegawa Y, Imaji A, Terabaru M, Nakamura R, Ohira N, Matsukura K, Seiwa K (2019) “Role of mycorrhizal associations in tree spatial distribution patterns based on size class in an old-growth forest.” Oecologia 189 (4):971–980

Shinohara N, Nakadai R, Suzuki Y, Terui A (2023) “Spatiotemporal dimensions of community assembly.” Popul Ecol 65 (1):5–16

Soininen J, McDonald R, Hillebrand H (2007) “The distance decay of similarity in ecological communities.” Ecography 30 (1):3–12

Soudzilovskaia NA, Vaessen S, Barcelo M, He J, Rahimlou S, Abarenkov K, Brundrett MC, Gomes SIF, Merckx V, Tedersoo L (2020) “FungalRoot: Global online database of plant mycorrhizal associations.” New Phytol 227 (3):955–966

Steidinger BS, Crowther TW, Liang J, Van Nuland ME, Werner GDA, Reich PB, Nabuurs GJ, de-Miguel S, Zhou M, Picard N, Herault B, Zhao X, Zhang C, Routh D, Peay KG, GFBI consortium (2019) “Climatic controls of decomposition drive the global biogeography of forest-tree symbioses.” Nature 569 (7756):404–408

Tuomisto H, Ruokolainen K, Yli-Halla M (2003) “Dispersal, environment, and floristic variation of western Amazonian forests.” Science 299 (5604):241–244

van der Putten WH, Bardgett RD, Bever JD, Bezemer TM, Casper BB, Fukami T, Kardol P, Klironomos JN, Kulmatiski A, Schweitzer JA, Suding KN, Van de Voorde TFJ, Wardle DA (2013) “Plant-soil feedbacks: The past, the present and future challenges.” J Ecol 101 (2):265–276

Viana DS, Chase JM (2019) “Spatial scale modulates the inference of metacommunity assembly processes.” Ecology 100 (2):e02576

Wickham H (2016) ggplot2: Elegant graphics for data analysis Springer-Verlag New York

Willig MR, Kaufman DM, Stevens RD (2003) “Latitudinal gradients of biodiversity: Pattern, process, scale, and synthesis.” Annu Rev Ecol Evol Syst 34 (1):273–309

Xing D, He F (2019) “Environmental filtering explains a U-shape latitudinal pattern in regional β-deviation for eastern North American trees.” Ecol Lett 22 (2):284–291

Yamawo A, Ohno M (2021). Joint evolution of mycorrhizal type, pollination, and seed dispersal mode in trees.” bioRxiv. doi.org/10.1101/2020.10.04.325282

Yamazaki D, Togashi S, Takeshima A, Sayama T (2020) “HIGH-RESOLUTION FLOW DIRECTION MAP OF JAPAN.” J Jpn Soc Civ Eng 8 (1):234–240 (in Japanese with English abstract)

Zhong, Y., Chu C, Myers JM, Gilbert GS, Lutz JA, Stillhard J, Zhu K, et al. (2021) “Arbuscular Mycorrhizal Trees Influence the Latitudinal Beta-Diversity Gradient of Tree Communities in Forests Worldwide.” Nat Commun 12 (1): 3137.

